# E2F1-3 activate Merkel cell polyomavirus early transcription and replication

**DOI:** 10.1101/2025.11.15.688648

**Authors:** Nicholas J.H. Salisbury, Supriya Patil-Amonkar, Ann Roman, Denise A. Galloway

**Affiliations:** Fred Hutchinson Cancer Center, Human Biology Division, Seattle, WA, 98109, USA; University of Washington, Department of Microbiology, Seattle, WA, 98195, USA

**Author notes:** **Author contributions: Nicholas Salisbury** – designed research, performed research, analyzed data, wrote the paper, reviewed and edited the paper. **Supriya Patil-Amonkar** – designed research, performed research, analyzed data, reviewed and edited the paper. **Ann Roman** – designed research, performed research, analyzed data, reviewed and edited the paper. **Denise Galloway** – designed research, analyzed data, reviewed and edited the paper.

**Keywords:** polyomavirus, E2F, Large Tumor antigen (LT), RB1

## Abstract

Merkel cell polyomavirus (MCPyV) is a DNA virus that establishes a persistent asymptomatic infection during childhood and can cause Merkel cell carcinoma (MCC) later in life. Its Large and Small Tumor antigens (LT, ST), splice variants of a common viral early transcript, drive viral replication and tumorigenesis by binding to and perturbing the function of host proteins. LT binds and inhibits RB1, deregulating E2F activity and host cell cycle control to permit viral replication during S phase. While the functions of LT and ST are relatively well characterized, how their expression is controlled is poorly defined. Here, we discovered that E2F1-3/DP1 dimers bind the MCPyV Non-Coding Control Region (NCCR) via a consensus E2 site close to the LT/ST transcriptional start site. Inhibiting E2F-NCCR binding, either by deleting the E2 site or treatment with a E2F small molecule inhibitor, downregulated LT/ST mRNA and protein expression in MCC cells and in 293A cells transfected with MCPyV. Our findings reveal an E2F/LT/RB1 positive feedback loop that appears to have evolved to support viral replication and is hijacked in MCC cells to promote cellular proliferation. Furthermore, we identified E2 sites in the NCCRs of PyVs closely related to MCPyV, including murine PyV, which mediate E2F/DP binding and potentiate viral early transcription. E2F/DP also bound weakly to the SV40 and BKPyV NCCRs, despite lacking an E2 site. Our findings challenge the prevailing model that PyV LT expression drives S phase entry and suggest, in contrast, that S phase entry stimulates PyV early transcription and replication.

**Significance statement:** Polyomaviruses (PyVs) express Large and Small Tumor antigens (LT, ST), splice variants of a common viral early transcript, that drive viral replication and tumorigenesis by binding and perturbing the function of host cells. LTs bind and inhibit RB1 via conserved LxCxE motifs, deregulating E2F activity and host cell cycle control to permit viral replication. Here, we discovered that E2F1-3 bind to the Non-Coding Control Regions (NCCRs) of many, but not all, PyVs to activate LT/ST transcription, revealing an E2F/LT/RB1 positive feedback loop that appears to have evolved to promote viral replication. Our findings challenge the existing paradigm that PyV LT expression drives S phase entry and suggest, in contrast, that S phase entry stimulates PyV LT/ST transcription and replication.

## Introduction

Polyomaviruses (PyVs) are circular, double-stranded DNA viruses with genomes of approximately 5 kilobases. They infect a wide range of vertebrates, including humans, and establish persistent infections that typically remain asymptomatic in immunocompetent hosts. To date, 16 human polyomaviruses have been identified(1). In immunosuppressed individuals, human PyVs can reactivate and cause severe disease, such as BKPyV-associated nephropathy and JCPyV-associated progressive multifocal leukoencephalopathy(2). Merkel cell polyomavirus (MCPyV) causes Merkel cell carcinoma (MCC), a deadly neuroendocrine skin cancer, and is the only human polyomavirus that definitively causes cancer. It was first identified integrated into the cellular genome of MCC tumors but has since been detected in healthy skin and other tissues(3, 4). Immunosuppressed individuals are at increased risk of developing PyV-associated diseases, although MCC develops in individuals with no overt immunodeficiency(5). While most individuals in the population are infected with PyVs during childhood(6, 7), it is unclear how these viruses maintain a persistent infection and what cellular cues prompt their reactivation.

PyV genomes are divided into three regions – early and late coding regions (ER, LR) and a non-coding control region (NCCR). ERs encode regulatory proteins that control PyV replication. The early proteins, or tumor antigens, of oncogenic polyomaviruses also having transforming potential(8–12). PyVs can be divided into three clades based on the structure of their ERs(13), with simian virus 40 (SV40), murine PyV, and MCPyV as archetype viruses for each clade. All PyVs express Large and Small Tumor antigens (LT, ST), which are splice variants of a common early transcript. The SV40 clade, which is the largest and includes all the human PyVs except trichodysplasia spinulosa-associated PyV (TSPyV) and MCPyV, express only these two early proteins. Members of the MuPyV clade, which includes TSPyV, express LT, ST and a Middle Tumor antigen (MT), which is a third splice variant of a common early transcript. MCPyV encodes LT and ST splice variants and a third protein, the Alternate LT Open reading frame (ALTO), which is encoded on the second exon of LT in +1 reading frame(13). LT/ST and ALTO are all expressed in cells infected with wt circular MCPyV but virus-positive MCC cells express only truncated variants of LT (LT-t) and wild-type ST(14, 15). ALTO is encoded in both the LT and ST transcripts but it is currently unknown how its expression is regulated, particularly how it is silenced in MCC when LT-t and ST are expressed.

LTs have two critical functions in viral replication. Firstly, they bind directly to the PyV origin of replication within the NCCR, unwind the viral DNA, and recruit host DNA replication machinery to the viral genome(16). Secondly, LTs bind to and inhibit RB1(17, 18), a negative regulator of the E2F family of transcription factors and host cell cycle progression. In normal quiescent cells, RB1 binds to E2F1-3 and their obligate partner proteins DP1/2, preventing them from activating expression of their target genes that mediate S-phase entry and DNA replication(19). In response to growth factor stimulation, cyclin-dependent kinases (CDKs) phosphorylate RB1, releasing it from E2F/DP and allowing E2Fs to activate transcription of their target genes(20). LTs bind to RB1 and disrupt the interaction between RB1 and E2F/DP, mimicking RB1 hyperphosphorylation. Thus, the current paradigm in the field is that PyV infection and subsequent expression of LT drives the host cell into S phase by activating E2F signaling. However, a recent study revealed that BKPyV can only express LT and replicate in cells already in S phase(21), yet how S phase signaling activates BKPyV LT expression is unknown.

PyV NCCRs contain the viral origin of replication, the transcriptional start sites of the early and late viral mRNAs and transcription factor (TF) binding sites that regulate early and late transcription. SV40’s NCCR has six SP1 binding sites, of which three are important for early transcription(22). SV40, along with BKPyV and JCPyV, also have NF-κB binding sites that regulate early transcription(23–25). The MuPyV NCCR contains binding sites for members of the AP-1 and ETS families of TFs(26). Little is known about the TFs that regulate MCPyV transcription either in MCC cells or during normal infection. TF binding sites have been predicted in the MCPyV NCCR but have not been experimentally validated(27, 28). NF-κB has been shown to bind and activate LT/ST transcription in MCC and 293 cells(29). However, we recently showed that MCPyV ALTO activates the NF-κB pathway in MCC cells and downregulating viral early transcription by binding to the MCPyV NCCR(15). Unlike JCPyV and BKPyV, MCPyV’s NCCR does not contain a consensus NF-κB binding site, so how NF-κB binds it is unclear. The MCPyV LT/ST transcriptional start site, which matches the consensus initiator element (Inr)(30), has been identified and is 23bp downstream of a consensus TATA box(31), which recruits TBP and RNA polymerase II to promoters (Fig. 1A). Here, we set out to identify TFs that bind the MCPyV NCCR and regulate viral early transcription in MCC cells and in 293A cells, the latter of which supports MCPyV LT/ST expression and replication.

**Figure 1.**
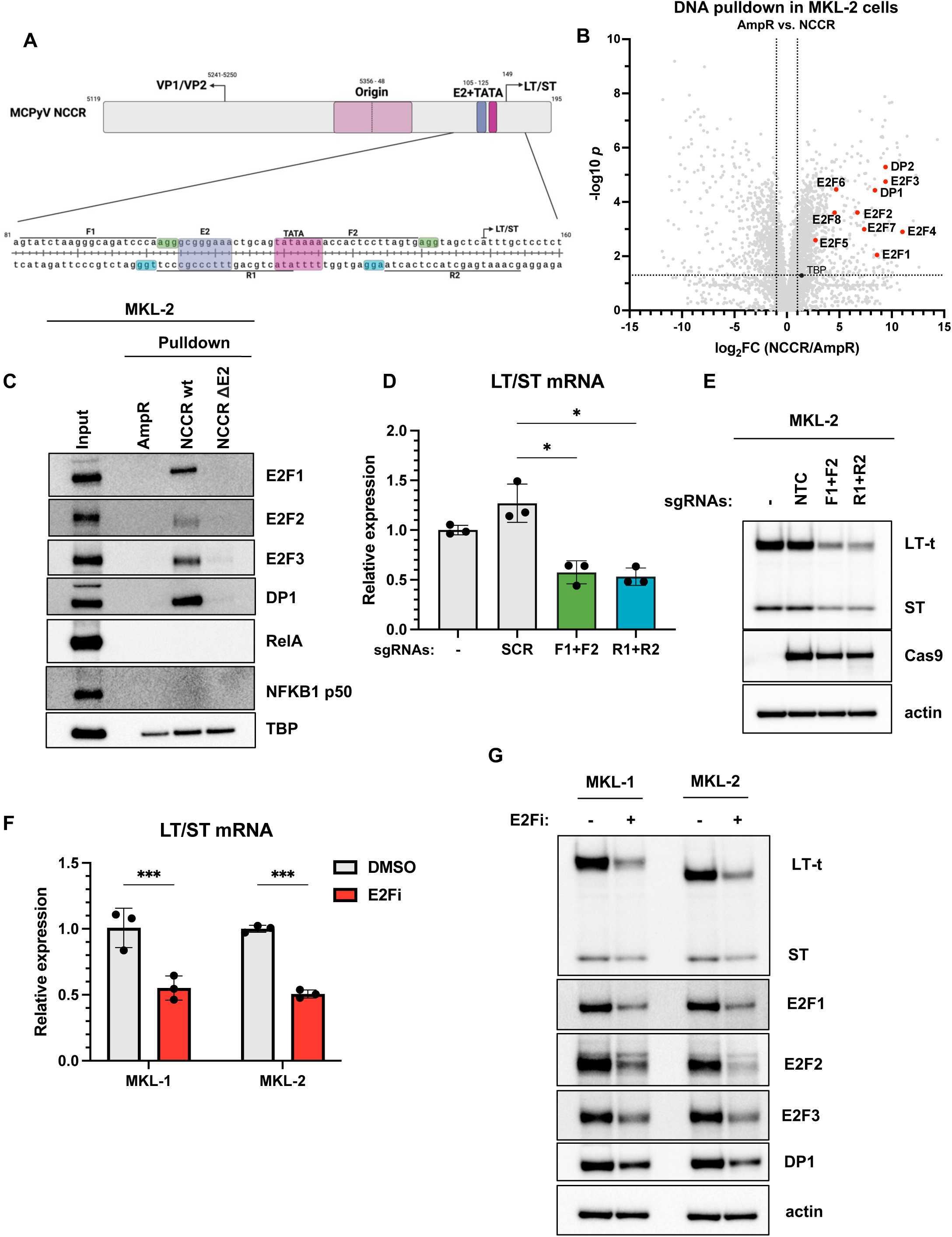
E2F1-3 bind the MCPyV NCCR and regulate viral early transcription in MCC cells. **A)** Schematic of the MCPyV NCCR highlighting the E2 site (purple), TATA box (pink) and PAM sites (green and blue) and sgRNA sequences (F1+F2, R1+R2) used in (*D)* and (*E)* to delete the E2 site and TATA box. **B)** Mass spectrometry analysis of DNA pulldown samples using MKL-2 cell lysates and AmpR and NCCR probes. **C)** Western blot analysis of proteins binding to AmpR and NCCR wt and ΔE2 probes in DNA pulldowns using MKL-2 cell lysates. **D)** qRT-PCR analysis of total LT/ST mRNA in MKL-2 cells transduced with lentiviral vectors encoding Cas9 and indicated sgRNAs and then harvested following 4 days of puromycin selection and 2 days of recovery. LT/ST mRNA expression is normalized by the expression of housekeeping gene 36B4. The experiment was performed with three biological replicates for each condition. Error bars represent the standard deviation of the mean. Statistical significance was calculated by one-way ANOVA test. * = p < 0.033. **E)** Western blots from cells analyzed in (*D)*. **F)** RT-qPCR analysis of total LT/ST mRNA expression in MKL-1 and MKL-2 cells treated with 40 μM HLM006474 or DMSO for 4 days. LT/ST mRNA expression is normalized by the expression of housekeeping gene 36B4. The experiment was performed with three biological replicates for each condition. Error bars represent the standard deviation of the mean. Statistical significance was calculated by two-way ANOVA test. *** = p < 0.0002. **G)** Western blots from cells analyzed in (*F*).

## Results

### E2F1-3 bind to the MCPyV NCCR and regulate viral early transcription in MCC cells

To identify TFs that bind to the MCPyV NCCR in MCC cells, we performed DNA pulldowns coupled with mass spectrometry using MKL-2 cell lysates and a dsDNA probe spanning the length of the MCPyV NCCR (464 bp, Fig. 1A&B). As a negative control for non-specific protein-DNA interactions, we created a dsDNA probe spanning the 3’ end of the ampicillin resistance gene (AmpR, 500 bp). Consistent with the presence of a TATA box, we observed enriched binding of TBP to the MCPyV NCCR relative to AmpR (FC = 2.64, *p* = 0.11). Contrary to the hypothesis that NF-κB activates LT/ST transcription in MCC cells, we did not detect binding of any NF-κB subunits (RelA, RelB, NFKB1, NFKB2) to the NCCR, nor AmpR. In total, we detected 980 proteins binding to the NCCR with FC > 2 and p < 0.05. Among the top hits, we detected binding of all eight members of the E2F family of transcription factors, E2F1-8, and their obligate dimerization partners, DP1/2, bound strongly to the NCCR (Fig. 1B). Since LT-t activates E2F1-3 signaling by inactivating RB1, we decided to focus our attention on examining the role on E2F1-3 in regulating LT/ST transcription. As validation of the proteomics, we examined the binding of E2F1-3, DP1, TBP, RelA and NFKB1 to the AmpR and NCCR probes by Western blot (Fig. 1C). Consistent with the mass spectrometry analysis, E2F1-3 and DP1 bound strongly to the NCCR probe but not to AmpR. TBP bound to both the AmpR and NCCR probes, but more strongly to NCCR than AmpR. While RelA and NFKB1 p50 are expressed in MKL-2 cells, we could not detect any binding of RelA and NFKB1 p50 to either probe by Western blot, consistent with our proteomic analysis. E2F/DP complexes bind to a highly conserved E2 site (5’-TTTSSCGC-3’) found in the promoters of its target genes(32). Analysis of the MCPyV NCCR revealed a consensus E2 site 6 bp upstream of the previously identified TATA box (Fig. 1A). To test whether E2F1-3 and DP1 bind to the NCCR via the E2 site, we generated an NCCR probe with the E2 site deleted (NCCR ΔE2). As predicted, E2F1-3 and DP1 did not bind to the NCCR ΔE2 probe (Fig. 1C). In summary, we conclude that E2F/DP complexes, but not NF-κB, bind to the MCPyV NCCR in MKL-2 cells under normal growth conditions.

To determine whether E2F binding to the MCPyV NCCR in MKL-2 cells regulates LT/ST transcription, we used CRISPR/Cas9 to delete the E2 site. We screened the NCCR for Protospacer Adjacent Motifs (PAM sites, 5’-NGG-3’) and identified two upstream of the E2 site and two downstream of the TATA box, allowing us to delete both the E2 site and TATA box but not the E2 site or TATA box alone (Fig. 1A). We delivered Cas9 and pairs of sgRNAs, one set annealing to 5’ strand (sgF1+F2) and another to 3’ strand (sgR1+R2), or a single non-targeting control sgRNA (sgNTC) via lentiviral vectors to MKL-2 cells. Consistent with E2F/DP and TBP binding to the NCCR to activate MCPyV transcription, we observed downregulation of LT/ST mRNA and proteins by ∼50% in MKL-2 sgF1+F2 and sgR1+R2 cells compared to either sgNTC or non-transduced control cells (Fig. 1D&E). These findings identify the E2 site and TATA box as critical promoter elements regulating LT/ST transcription in MKL-2 cells.

As further validation of E2F1-3 regulating LT/ST transcription in MCC cells, we treated MKL-1 and MKL-2 cells with a small molecule E2F inhibitor (HLM006474) and assessed its effects on LT/ST mRNA and protein expression (Fig. 1F&G). As previously observed(33), HLM006474 treatment significantly downregulated E2F1-3 and DP1 protein levels in both MKL-1 and MKL-2 cells. Consistent with E2F1-3 activating LT/ST transcription, HLM006474 treatment downregulated LT/ST mRNA and protein levels in both cell lines. These observations identify E2F1-3 as critical activators of LT/ST transcription in MCC cells.

### E2F1-3 bind to the MCPyV NCCR and regulate viral early transcription and replication in 293A cells

Next, we sought to identify TFs that regulate LT/ST transcription during MCPyV infection. Although dermal fibroblasts are thought to be the natural MCPyV host cell(34), 293 cells have been shown to support MCPyV LT/ST expression and viral replication(35). 293 cells and 293 derivatives, such as 293A cells, all express adenovirus 5 E1A, which binds and inhibits RB1 via its LxCxE motif(36, 37), in an analogous manner to PyV LTs. Thus, E2F1-3 are constitutively active in 293 cells. We performed DNA pulldowns in 293A cells and detected binding of E2F and DP family members to the MCPyV NCCR by mass spectrometry and Western blot (Fig. 2A&B). As in MKL-2 cells, E2F1-3 and DP1 binding to the NCCR probe is abolished when the E2 site is deleted. In 293A cells, TBP was not detected by mass spectrometry, but we observed a similar pattern of binding to the AmpR and NCCR probes as observed in MKL-2 cells by Western blot. We also analyzed the binding of NF-κB subunits to the NCCR in 293A cells. We did not detect any binding of RelA or RelB to either the AmpR or NCCR probe via mass spectrometry and validated lack of RelA binding by Western blot. We could detect some binding of NFKB1 and NFKB2 to the NCCR probe by mass spectrometry but significantly weaker than E2F/DP binding. We analyzed NFKB1 binding by Western blot and were able to detect a very faint band in the NCCR wt and ΔE2 pulldowns. We conclude that E2F/DP dimers bind significantly stronger to the MCPyV NCCR than NF-κB in 293A cells under normal growth conditions.

**Figure 2.**
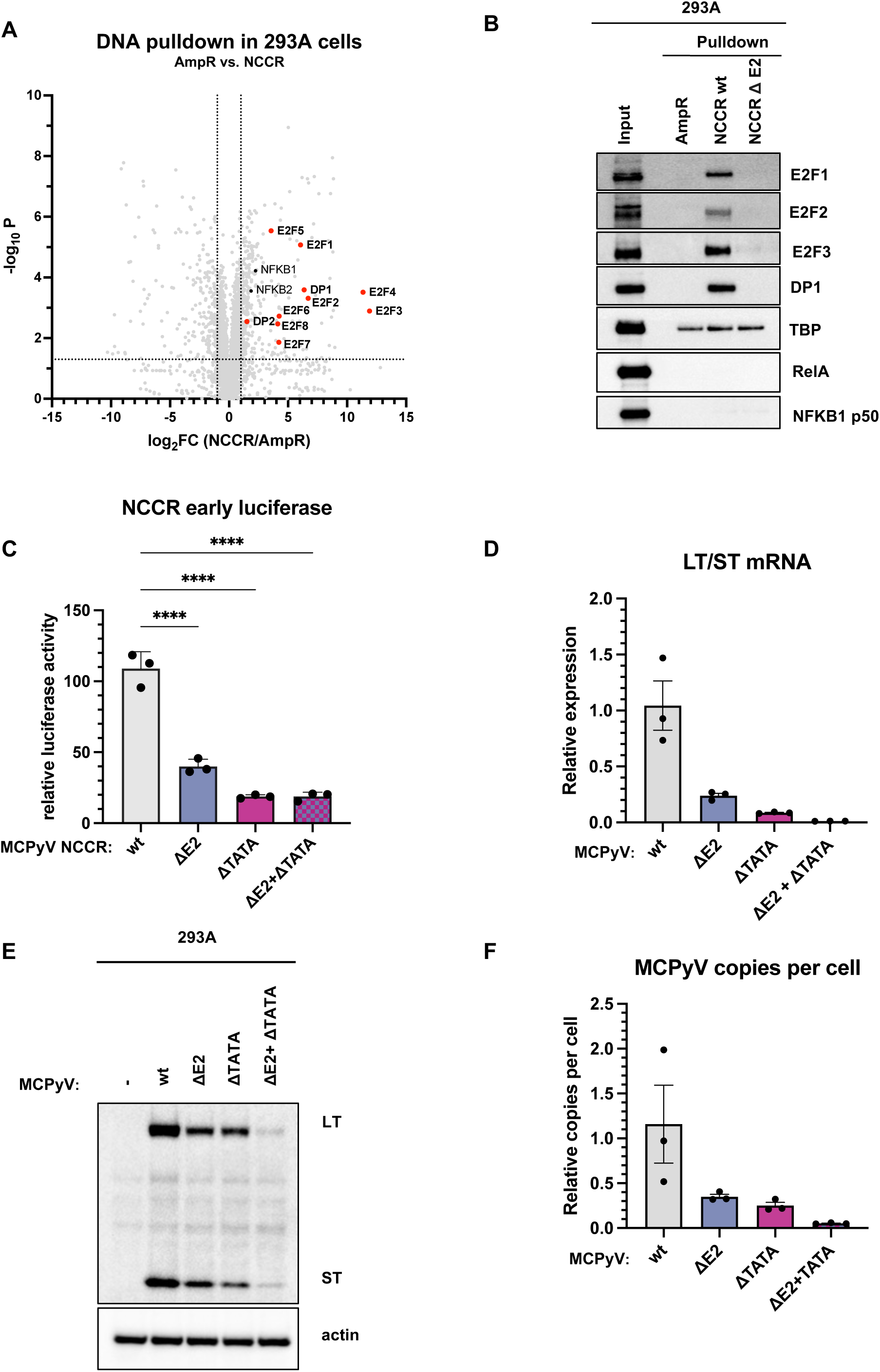
E2F1-3 bind the MCPyV NCCR and regulate LT/ST transcription and viral replication in 293A cells. **A)** Mass spectrometry analysis of DNA pulldown samples using 293A cell lysates and AmpR and NCCR probes. **B)** Western blot analysis of proteins binding to AmpR and NCCR wt and ΔE2 probes in DNA pulldowns using 293A cell lysates. **C)** Relative luciferase activity in 293A cells transfected with MCPyV early firefly luciferase and CMV renilla luciferase plasmids. Firefly luciferase activity was normalized by renilla luciferase activity. The reported luciferase activity is relative to cells transfected with an empty vector firefly luciferase reporter and CMV renilla plasmid. The experiment was performed with three biological replicates for each condition. Error bars represent the standard deviation of the mean. Statistical significance was calculated by one-way ANOVA. * p < 0.033, ** p < 0.002. **D)** RT-qPCR analysis of total LT/ST mRNA in 293A cells transfected with MCPyV variants and harvested four days later. LT/ST mRNA expression is normalized by the expression of housekeeping gene 36B4. The experiment was performed with three biological replicates for each condition. Error bars represent the standard deviation of the mean. Statistical significance was calculated by one-way ANOVA but the p-values for the comparisons between MCPyV wt and any of the variants were all greater than 0.033 due to the high variation in MCPyV wt. **E)** Western blots from cells analyzed in (*D)*. Blots have been performed with for three biological replicates and one set of representative blots are shown. **F)** qPCR analysis of relative MCPyV copy number in 293A cells analyzed in *(D)* and *(E)*. The experiment was performed with three biological replicates for each condition. Error bars represent the standard deviation of the mean. Statistical significance was calculated by one-way ANOVA but the p-values for the comparisons between MCPyV wt and any of the variants were all greater than 0.033 due to the high variation in MCPyV wt.

As a first step to determine whether E2Fs regulate MCPyV LT/ST transcription in 293A cells, we measured the effect of deleting the E2 site from the NCCR on viral early transcription using our previously described MCPyV NCCR early luciferase vector(15). Loss of the E2 site (ΔE2) caused a 60% reduction in luciferase activity (Fig. 2C). Deleting the TATA box (ΔTATA) caused an 80% reduction in luciferase activity. Loss of both the E2 site and TATA box (ΔE2+ΔTATA) did not significantly reduce luciferase activity below that of the ΔTATA reporter. In all cases, the luciferase activity of the mutant NCCR reporters was higher than that of a negative control luciferase reporter that contained no promoter sequence inserted upstream of the luciferase coding sequence. Next, we deleted the E2 site, TATA box or both in a recombinant MCPyV minicircle clone(38) and assessed the ability of the mutant MCPyV genomes to express LT/ST and replicate in 293A cells (Fig 2D-F). Deletion of the E2 site reduced LT/ST mRNA by 76% at four days post transfection. Deletion of the TATA box reduced LT/ST mRNA by 91% and deleting both caused a 99% reduction. We observed a similar trend in LT and ST protein expression at the same time point (Fig 2E). Consistent with decreased LT/ST expression, MCPyV ΔE2 had a 70% reduction in copies per cell compared to the wild type at four days post-transfection. MCPyV ΔTATA genome copies were reduced by 80% at the same time point, and MCPyV ΔE2+ΔTATA was reduced by 95%. Overall, our findings demonstrate that the E2 site and TATA box are critical promoter elements that control LT/ST transcription in 293A cells, as observed in MKL-2 cells, and indicate that E2F1-3 are critical activators of MCPyV replication.

### A subset of PyV NCCRs contain E2F binding sites that regulate PyV early transcription

A recent report revealed that BKPyV can only express LT and replicate in cells already in S phase(21), but the mechanism controlling BKPyV LT expression in S phase is currently unknown. Given the well-established role of E2Fs in regulating S phase entry and our findings so far, we postulated that E2Fs might bind the NCCRs of BKPyV and other PyVs to regulate LT/ST expression. We searched for E2 sites, TATA boxes and Inr elements in the NCCRs from a range of nine mammalian polyomaviruses, including three members each of the MCPyV, MuPyV, and SV40 clades (Fig. 3A). In the NCCR of Gorilla PyV, which is closely related to MCPyV, we identified a consensus E2 site (5’-TTTSSCGC-3’), TATA box and Inr element in a very similar arrangement to the MCPyV NCCR. The NCCR of otomops PyV, more distantly related to MCPyV, contains a consensus TATA box and a variant E2 site (5’-CTTSSCGC-3’) but no Inr element. Structural and biochemical studies have shown that that E2F/DP complexes can tolerate mismatches in the TTT part of the E2 site motif, while the central SSCG part of the motif is critical(32, 39). Murine polyomavirus also contains a variant E2 site (5’-CTTSSCGC-3’), which has not been previously identified as far as we are aware and is just downstream of the previously identified TATA box(40, 41). MuPyV also contains a consensus Inr element downstream of the TATA box and putative E2 site. While closely related to MuPyV, hamster PyV and TSPyV’s NCCRs show a different structure with consensus E2 sites close to the start codon of their respective VP2s. In contrast to our initial prediction, BKPyV’s NCCR does not contain a consensus or variant E2 site. The NCCRs of JCPyV and SV40 also do not. In conclusion, E2 sites are not a universal feature of PyV NCCRs but are found in the NCCRs of MT- and ALTO-encoding PyVs.

**Figure 3.**
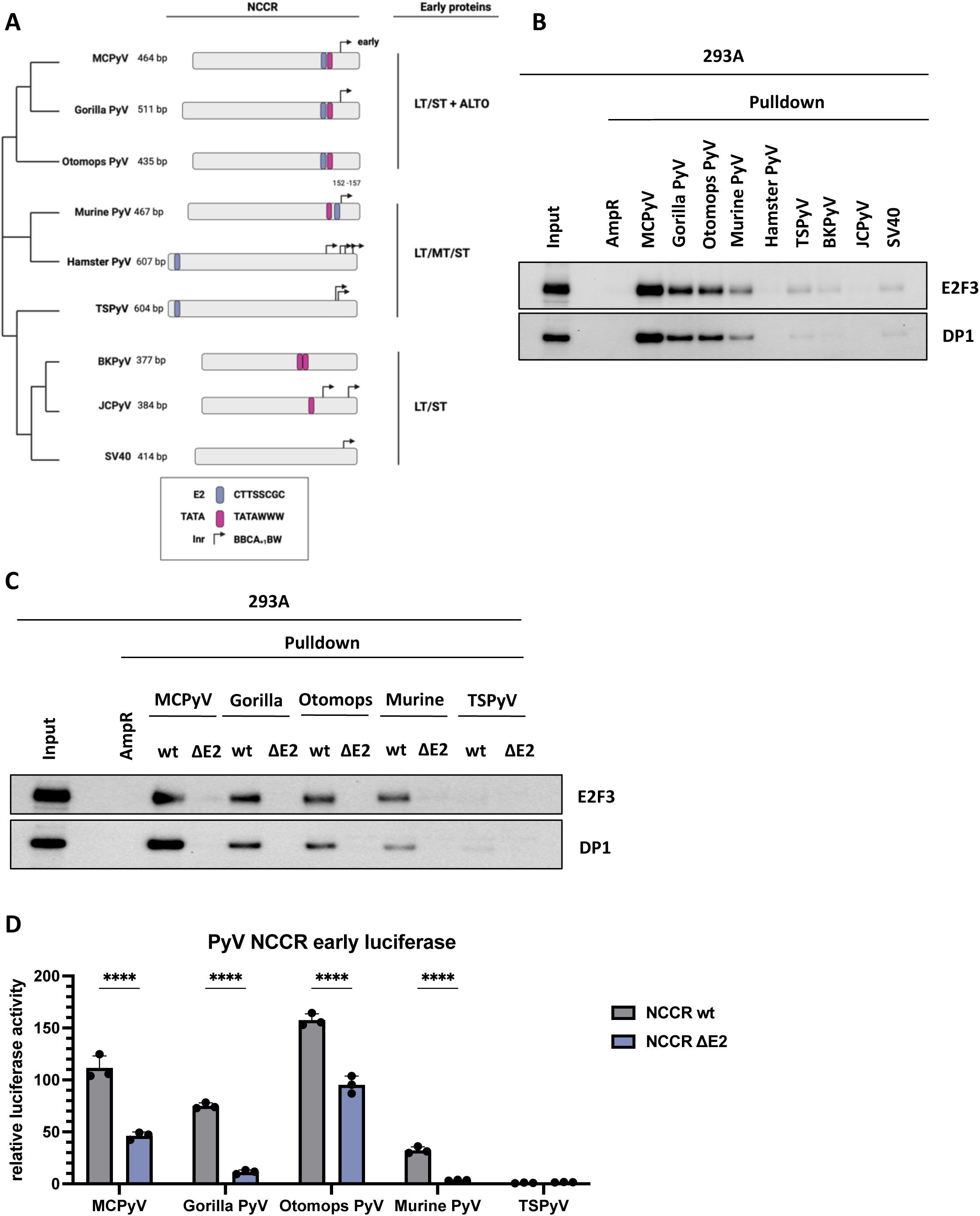
A subset of PyV NCCRs contain E2 sites that regulate early transcription. **A)** Schematic of PyV NCCRs highlighting known and putative promoter elements including E2 sites, TATA boxes, and initiator (Inr) elements. **B)** Western blots of DNA pulldown samples using PyV NCCRs as bait and 293A cells lysates as prey. **C)** Western blots of DNA pulldown samples using wild type PyV NCCRs and variants with the putative E2 sites deleted (ΔE2) as bait and 293A cell lysates as prey. **D)** Relative luciferase activity in 293A cells transfected with PyV early firefly luciferase and CMV renilla luciferase plasmids. Firefly luciferase activity was normalized by renilla luciferase activity. The reported luciferase activity is relative to cells transfected with an empty vector firefly luciferase plasmid and CMV renilla luciferase plasmid. The experiment was performed with three biological replicates for each condition. Error bars represent the standard deviation of the mean. Statistical significance was calculated by two-way ANOVA. **** p < 0.0001.

Next, we examined the ability of E2F3 and DP1 from 293A cells to bind to PyV NCCRs in a DNA pulldown (Fig. 3B). As observed previously, E2F3 and DP1 both bound strongly to MCPyV’s NCCR but not to the negative control AmpR probe. As predicted based on the presence of consensus or variant E2 sites, E2F3 and DP1 bound to the NCCRs of gorilla, otomops and murine PyVs and weakly to TSPyV’s. We did not detect any E2F3/DP1 binding to hamster PyV’s NCCR, despite the presence of a consensus E2 site. As predicted, deleting the E2 site in the NCCRs of gorilla, otomops, murine and TSPyV abolished E2F3/DP1 binding (Fig. 3C). While lacking an E2 site, we detected weak E2F3/DP1 binding to the NCCRs of BKPyV and SV40 but not to JCPyV (Fig. 3B). In conclusion, E2F3/DP1 bind to many but not all PyV NCCRs, which is frequently but not always mediated by a consensus or variant E2 site.

Finally, to determine whether E2F/DP binding to PyV NCCRs regulates early transcription, we generated PyV early luciferase reporters with either wild-type or ΔE2 NCCRs and compared their luciferase activities when transfected in 293A cells (Fig 3D). As observed previously, deleting the E2 site in MCPyV’s NCCR reduced luciferase activity by 60%. Deleting the E2 site in gorilla PyV NCCR reduced luciferase activity by 85%, by 40% in otomops PyV’s NCCR, and by 90% in MuPyV’s NCCR. Our TSPyV early luciferase reporter showed no activity above the level of the empty vector negative control and no change following deletion of the E2 site, suggesting that E2Fs do not activate TSPyV early transcription. In conclusion, E2F/DP complexes bind to the NCCRs of gorilla, otomops and murine PyV and activate LT/ST transcription.

## Discussion

PyV early proteins play critical roles in viral replication and the etiology of PyV-associated diseases, including tumorigenesis. Identifying host transcription factors that regulate viral transcription is critical for understanding how PyVs maintain a persistent infection throughout the life of the host and for identifying therapeutic strategies to treat PyV-associated diseases. PyV NCCRs contain viral transcriptional start sites and TF binding sites. While some host TFs have been identified that bind to the NCCRs for a handful of PyVs, we are still far from a complete picture of how PyV transcription is regulated. In particular, very little is known about the host transcription factors that regulate MCPyV early transcription during normal infection and in MCC cells. A previous study identified NF-κB as an activator of LT/ST transcription in MCC and 293 cells(29). The authors described a model whereby NF-κB binds to the NCCR and activates LT/ST transcription when expressed at low levels and inhibits LT/ST transcription when the pathway is “overstimulated” or RelA is overexpressed. They demonstrated that RelA binds to the NCCR in a DNA pulldown when overexpressed, but not at endogenous levels. We recently showed that MCPyV ALTO activates NF-κB signaling in both MCC and 293A cells. In our hands, ALTO or overexpressed RelA downregulated endogenous LT/ST transcription in MCC cells and an MCPyV early luciferase reporter in 293A cells(15). We detected binding of endogenous NF-κB to the MCPyV NCCR via a DNA pulldown assay using MKL-2 cells expressing ALTO. Here, we observed that NF-κB subunits do not bind to the MCPyV NCCR in MKL-2 cells in the absence of ALTO, indicating that NF-κB does not activate LT/ST transcription in MCC cells. We detected very weak binding of NFKB1 and NFKB2 to the NCCR in 293A cells in the absence of ALTO. Since NFKB1 and NFKB2 do not contain transcriptional activation domains, we predict that NFKB1 and NFKB2 do not activate MCPyV early transcription in 293A cells. While NF-κB binding sites have been predicted in the MCPyV NCCR, the predicted sites do not match the consensus κB sequence (5’-GGGRNYYYCC-3’) that is found in the BKPyV and JCPyV NCCRs(27, 28). It remains unclear how NF-κB, once activated by ALTO or overexpressed, binds to the MCPyV NCCR in the absence of a consensus κB site, and how it downregulates LT/ST transcription.

In the present study, we identified E2F1-3 as critical activators of LT/ST transcription and demonstrated that they bind the MCPyV NCCR via a consensus E2 site, just upstream of a previously reported consensus TATA box and a consensus initiator element (Inr). These three elements appear to form the core promoter regulating LT/ST transcription. As far as we are aware, ours is the first study identifying E2Fs as critical regulators of PyV early transcription. Our findings reveal a E2F/LT/RB1 positive feedback loop whereby E2F1-3 activate expression of LT, which binds to and inhibits RB1, sustaining E2F activity (Fig. 4A). This positive feedback loop likely evolved to promote viral replication but is hijacked in MCC cells to promote cellular proliferation. The same E2F/LT/RB1 feedback loop appears to be present in both oncogenic (MCPyV, MuPyV, SV40) and non-oncogenic PyVs (BKPyV). Our findings also explain why 293 cell lines support robust LT/ST expression as they express adenovirus 5 E1A, which binds and inactivates RB1 in a similar manner to PyV LTs(42). Thus, 293(A) cells have constitutively active E2F signaling and are primed for MCPyV early transcription (Fig. 4B). Identifying E2Fs as critical regulators of MCPyV LT/ST expression in MCC cells opens the possibility of using E2F inhibitors to treat MCC. Two previous studies identified small molecules that downregulate LT/ST transcription by inhibiting the histone acetyltransferases (HATs) CREBBP and EP300(29, 43). CREBBP and EP300 have been shown to bind the transcriptional activation domain of E2F1(44, 45), suggesting that E2Fs recruit CBP and EP300 to the MCPyV NCCR to acetylate NCCR-associated histones and drive LT/ST transcription. Future work ought to assess whether treatment of MCC cells with E2F and CREBBP/EP300 inhibitors selectively kill MCC cells compared to normal proliferating cells.

**Figure 4.**
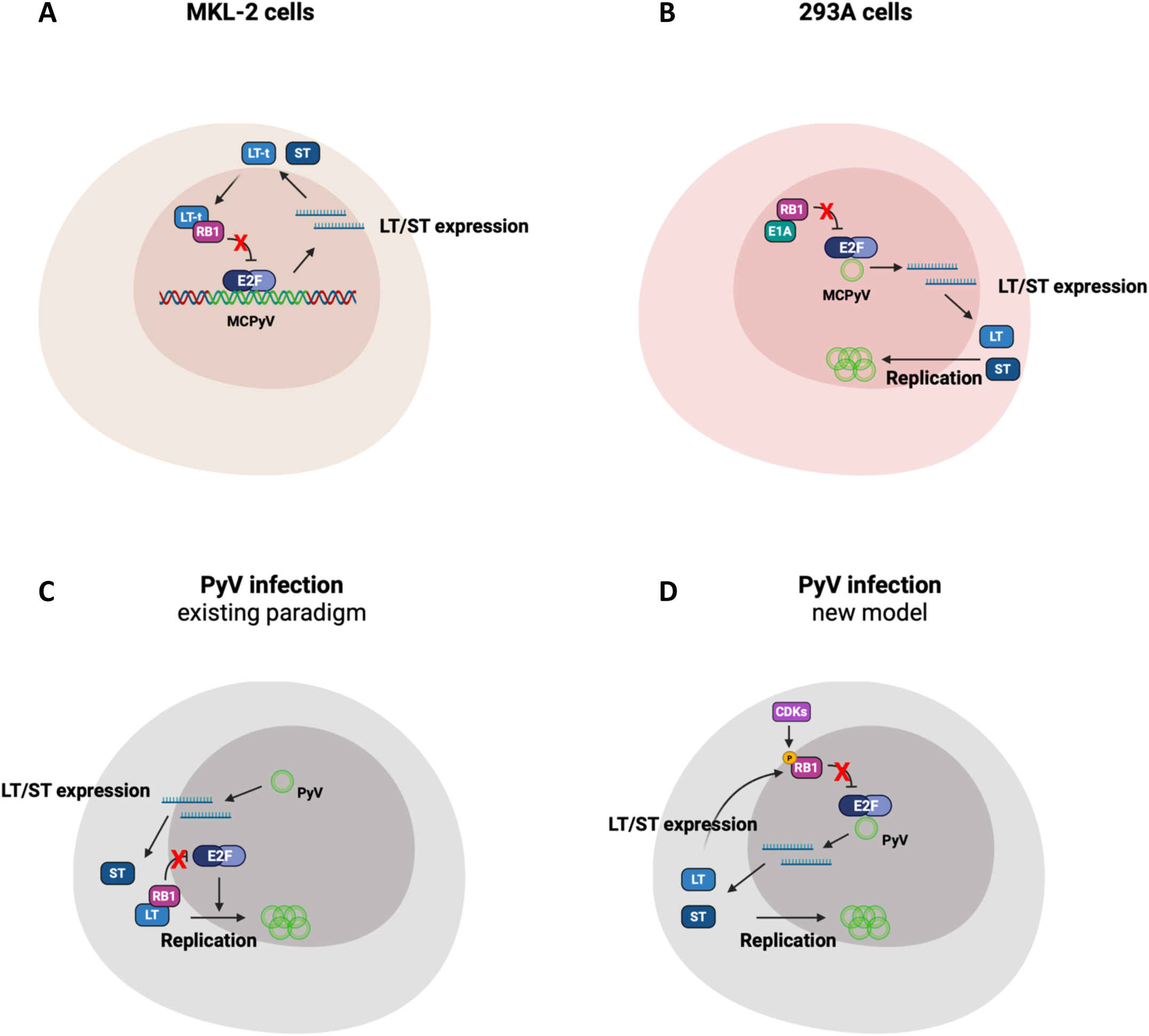
E2F signaling activates PyV LT/ST expression and viral replication. **A)** In MKL-2 cells, a mutant MCPyV genome that encodes truncated LT (LT-t) is integrated in the host cellular genome. LT-t binds and inhibits RB1, activating E2F1-3, via its LxCxE motif. E2F/DP dimers bind to the E2 site in the MCPyV NCCR and drive LT/ST transcription. Together, LT-t/RB1/E2F signaling creates a positive feedback loop that deregulates E2F activity, sustains LT/ST expression, and promotes tumorigenesis. **B)** In 293A cells, adenovirus 5 E1A is expressed, which binds and inhibits RB1 via its LxCxE motif and activates E2F signaling. Thus, 293A cells are primed for MCPyV LT/ST expression and replication. **C)** Expression of PyV LTs in quiescent cells stimulates cell proliferation by binding to RB1 and activating E2F signaling. Hence, it was previously thought that PyV infections stimulate host cell proliferation, with E2Fs contributing to viral replication by upregulating expression of their target genes. **D)** Our findings suggest that host cell entry into S phase, with RB1 inactivated by cyclin-dependent kinases (CDKs) and E2Fs active, activates LT/ST expression and viral replication. Once expressed, LT can bind and maintain RB1 inactivation, prolonging S phase to promote PyV replication.

Our results suggest that productive MCPyV infection and replication are intimately tied to the host cell cycle. Dermal fibroblasts have been shown to support MCPyV entry, gene expression and replication *in vitro*(34). Fibroblasts must be serum-starved for MCPyV entry and then serum-stimulated for robust LT expression and viral replication. Our results help explain why serum stimulation, a known activator of E2F signaling, is critical for MCPyV LT expression and replication. In addition to LT expression, E2F signaling might also indirectly regulate MCPyV nuclear entry. In contrast to other PyVs, MCPyV does not enter the nucleus via the nuclear pore complex but, rather, takes advantage of cell-cycle dependent nuclear envelope breakdown to gain access(46). Taking all our findings together, it appears that, following MCPyV endocytosis, an infected cell appears to need to pass through mitosis for MCPyV to traffic to the nucleus and then pass through the G_1_-S checkpoint again before MCPyV can express its genes and begin replicating.

Currently, the virus-positive MCC cell of origin is unknown. Evidence points to MCC arising from an epithelial cell type or possibly a cell of neuronal lineage, although fibroblasts and pre-/pro-B cells have also been proposed(47–50). LT mutation and clonal integration of the mutant MCPyV genome appear to be early events in MCC tumorigenesis(14). As LT/ST expression appears to depend on the host cell entering S phase, our findings point to the MCC cell of origin being a stem or progenitor cell type, perhaps a basal keratinocyte or a Merkel cell progenitor. In MKL-2 cells, deletion of the E2 site and TATA box significantly downregulated, but did not completely abolish, LT/ST transcription and protein expression (Fig. 1D&E), indicating that other host TFs bind the NCCR in MCC cells and contribute to LT/ST transcription. It is plausible that these factors could be cell lineage transcription factors. MCPyV infection is thought to be relatively promiscuous with MCPyV pseudovirions carrying a CMV-GFP reporter plasmid able to enter a wide variety of different cell types and express GFP(51). In another study, MCPyV particles were shown to infect fibroblasts and keratinocytes readily and Merkel cells at very low frequencies (1 in 122 KRT20+ cells), but LT and VP1 expression was only detected in fibroblasts(34). These data suggest that the NCCR could be a key determinant of viral tropism. Since E2Fs are essential genes expressed in all cell types, it is reasonable to suspect that mesenchymal lineage TFs bind the NCCR to facilitate MCPyV gene expression in fibroblasts. Further work will be needed to address what these TFs in fibroblasts are and whether these same TFs are expressed and contribute to LT/ST transcription in MCC cells or if other cell lineage TFs plays a role, which could provide insight into the MCC cell of origin.

As a productive BKPyV infection appears to require the host cell to be actively replicating(21), we examined E2F/DP binding across a representative collection of PyVs. We identified E2 sites that regulate early transcription in gorilla, otomops, and murine PyVs that are all relatively closely related to MCPyV. We detected very weak binding of E2F/DP to an E2 site in the NCCR of TSPyV, located close to the VP2 start codon, however, a TSPyV early luciferase reporter was not active in 293A cells, suggesting that this TSPyV E2 site does not regulate early transcription. Future work will be required to assess whether E2Fs activate TSPyV late transcription. While we could not identify an E2 site in SV40 or an archetype strain of BKPyV (NEB-10), we did detect weak binding of E2F/DP to their NCCRs. It is therefore plausible that E2F1-3 activate BKPyV early transcription, explaining why BKPyV only expresses LT in host cells already in S phase. Further work will be required to examine how E2F/DP dimers bind to the SV40 and BKPyV NCCRs and determine if E2Fs activate SV40 and BKPyV LT/ST transcription and replication.

Overall, our findings support a model whereby PyVs exist in a dormant phase with little to no gene expression if the host cell is in G_0_ or G_1_ phases of the cell cycle and begin to express their early genes and replicate when the host cell enters S phase. Previously, it was thought that PyV infection and LT expression stimulates host cell S phase entry (Fig. 4C). Our study of MCPyV and previous findings in BKPyV suggest, in fact, that host cell S phase entry drives PyV replication (Fig. 4D). Once expressed, LT can bind to RB1 and sustain E2F activity, prolonging S phase and providing more time for unlicensed PyV replication. As most PyV infections are asymptomatic, we propose that PyVs spend most of their time in a dormant state in non-dividing host cells. However, if a PyV-infected cell receives a signal to divide, perhaps during a wound healing response following tissue damage, it will also stimulate PyV replication, allowing the virus to spread to other cells and maintain a persistent infection in the regenerating tissue. Furthermore, in a dormant state with little to no viral gene expression, PyVs will be able to evade host immune detection and persist for the life of the host.

## Materials and Methods

### Cell lines

MCC cell lines, MKL-1 and MKL-2, were previously obtained from the Chang & Moore lab (U. Pittsburgh) and were maintained in RPMI 1640 HEPES medium (Gibco) supplemented with 10% fetal bovine serum (FBS, Corning), and penicillin-streptomycin solution (Gibco). 293A cells (Invitrogen) were cultured in Dulbecco’s modified Eagle’s medium (Gibco) supplemented with 10% FBS, glutaMAX (Gibco), non-essential amino acids (Gibco) and penicillin-streptomycin solution. All cells were cultured at 37°C with 5% CO_2_ in a humidified atmosphere.

### Plasmids and subcloning

MCPyV minicircle genomic DNA was prepared from pMC.BESPX-MCV-HF (kindly provided by the Chang and Moore lab) as previously described(38). MCPyV minicircle variants were produced using the Q5 site-directed mutagenesis kit (NEB) following manufacturer’s instructions. PyV NCCR variants were synthesized (IDT) and cloned into pGL4.11 (Promega) using InFusion HD cloning kit (Takara) following manufacturer’s instructions. CMV-renilla plasmid pGL4.75 was obtained from Promega. Lentiviral Cas9+sgRNA vectors were synthesized by VectorBuilder. See Supplementary Table 1 for sgRNA sequences.

### DNA pulldown assays

Small scale DNA pulldowns were performed as described previously(15), using 5 μg biotinylated dsDNA probe, 50 μL streptavidin magnetic beads, and 500 μg cell lysate in 500 μL. Following PCR amplification and purification, dsDNA probes were Sanger sequenced to verify sequences. See Supplementary Table 2 for the sequences of 5’-biotinylated PCR primers used to generate AmpR and PyV NCCR dsDNA probes and Supplementary Table 3 for the sequences of the dsDNA probes.

### Mass spectrometry sample prep and analysis

For large scale DNA pulldowns for mass spectrometry analysis, 30 μg biotinylated dsDNA probe, 300 μL streptavidin magnetic beads, and 3 mg MKL-2 or 293A cell lysate was used per replicate. Following washing the beads five times in 20 mM Tris-HCl pH 7.5, 50 mM KCl to remove detergents, the magnetic streptavidin beads and bound proteins were resuspended in 8 M urea in 100 mM Tris pH 8.5. Protein disulfide bonds were reduced with 10 mM tris (2-carboxyethyl) phosphine with mixing at room temperature for 20 min. The reduced proteins were alkylated using 20 mM 2-chloroacetamide with mixing in the dark at room temperature for 30 min. Samples were digested with rLys-C protease (500 ng) with mixing at 37°C for 4 hr. The concentration of urea was reduced to 1 M with 100 mM Tris pH 8.5 and 10 mM calcium chloride was added. Trypsin protease (500 ng) was added and the reaction was mixed overnight at 37°C. Samples were removed from heat, spun, put on a magnetic rack and the supernatant was collected. The beads were washed with 0.1% trifluoroacetic acid and the solutions were combined and dried on a speedvac. Samples were desalted over TopTip (LC Packings), eluted with 15 mM ammonium formate pH 2.8 and dried on a speedvac. The desalted samples were brought up in 2% acetonitrile in 0.1% formic acid prior mass spectrometry analysis.

Samples were analyzed by LC/ESI MS/MS with a Thermo Scientific Vanquish Neo UHPLC (Thermo Scientific, Waltham, MA) coupled to a tribrid Orbitrap Ascend with FAIMS pro (Thermo Scientific, Waltham, MA) mass spectrometer. In-line desalting was accomplished using a reversed-phase PepMap Neo Trap Cartridge C_18_ (300 μm × 5 mm, 5-μm, 100 Å C_18_ resin; Thermo Scientific, Waltham, MA) followed by peptide separations on a reversed-phase column (75 μm × 270 mm) packed with ReproSil-Pur C_18_AQ (3-μm 120 Å resin; Dr. Maisch, Baden-Würtemburg, Germany) directly mounted on the electrospray ion source. A 90-minute gradient from 8% to 30% B (80% acetonitrile in 0.1% formic acid/water) at a flow rate of 300 nL/minute was used for chromatographic separations. LC/MS/MS data were collected using data dependent acquisition methods with the MS survey scans detected in the Orbitrap. MS/MS spectra were detected in the linear ion trap using HCD activation. Selected ions were dynamically excluded for 60 seconds after a repeat count of 1.

Data analysis was performed using Proteome Discoverer 3.1 (Thermo Scientific, San Jose, CA). The data were searched against Uniprot Human (UP000005640_Human_2023_11_09.fasta), Merkel Cell (NCBI_MerkelCellPolyomavirus_040824.fasta) and common contaminants (cRAPome). Searches were performed with settings for the proteolytic enzyme trypsin. Maximum missed cleavages were set to 2. The precursor ion tolerance was set to 10 ppm and the fragment ion tolerance was set to 0.6 Da. Dynamic peptide modifications included oxidation (+15.995 Da on M). Dynamic modifications on the protein terminus included acetyl (+42.011 Da on N-terminus), Met-loss (-131.040 Da on M) and Met-loss+Acetyl (-89.030 Da on M) and static modification carbamidomethyl (+57.021 on C). Minora was used for peak abundance and retention time alignment. Sequest HT was used for database searching. All search results were run through Percolator for scoring. Raw quantitative results were transformed to log_2_ scale and normalized to the median value across samples. Missing data were imputed with half of the global minimum value. P-values for pairwise comparisons were calculated by t-test. Mass spectrometry data sets are available in the Supporting Information dataset file and relevant legends in the Supporting Information Appendix.

### Lentiviral transduction

To generate lentiviral particles, 293T cells were transfected with psPAX2 and pMD2.G (gifts from Didier Trono – Addgene #12260, #12259) and Cas9+sgRNA lentiviral vectors. Supernatants containing lentiviral particles were harvested 48 h post transfection, 0.45 μM filtered, and then supplemented with 6 μg/mL polybrene. MKL-2 cells were incubated with lentiviral supernatants for 24 h followed by 24 h recovery in fresh growth medium. Transduced cells were selected with 0.5 μg/mL puromycin for 4 days and then allowed to recover for 2 days in fresh growth medium lacking puromycin.

### RNA extraction and quantitation by RT-qPCR

Total RNA was extracted from cells using RNeasy mini kit (Qiagen) following manufacturer’s instruction including DNase I on-column digestion to remove gDNA. cDNA synthesis was performed using SuperScript VILO cDNA synthesis kit (Invitrogen) following manufacturer’s instructions. qPCR reactions were performed using PowerUP SYBR Green master mix (Applied Biosystems) and a StepOnePlus real-time qPCR system (Applied Biosystems). qPCR reactions contained 300 nM primers and were run on the fast cycling setting. See Supplementary Table 4 for qPCR primer sequences.

### Protein extraction, Western blotting and antibodies

Proteins were extracted and Western blotting performed as previously described(15). The following antibodies were used: E2F1 (#3742, Cell Signaling Technologies), E2F2 (ab31895, Abcam), E2F3 (ab320731, Abcam), DP1 (ab124678, Abcam), TBP (#44059, Cell Signaling Technologies), RelA (#8242, Cell Signaling Technologies), NFKB1 (#12540, Cell Signaling Technologies), Ab5 for MCPyV LT and ST (a kind gift from the DeCaprio lab, Dana-Farber Cancer Institute), Cas9 (#14697, Cell Signaling Technologies), actin-HRP (#5125, Cell Signaling Technologies).

### MCPyV genome transfection

293A cells were seeded at 2.5x10^6^ cells per 10 cm plate. The following day, cells were transfected with 5 μg of MCPyV minicircle DNA using TransIT 293 (Mirus Bio) following manufacturer’s instructions. Cells were harvested 4 days post-transfection for further analysis.

### gDNA extraction and quantitation by qPCR

gDNA was extracted from 293A cells using PureLink genomic DNA mini kit following manufacturer’s instruction. Typically, 2.5x10^6^ cells were used for each extraction. qPCR reactions were performed using SYBR Green PowerUP master mix (Applied Biosystems), a StepOnePlus real-time qPCR system (Applied Biosystems) and 10 ng of gDNA as the input for each qPCR. See supplementary data for details of qPCR primers, which are identical to primers described previously(38).

### Luciferase assays

Luciferase assays were performed using MCPyV NCCR early luciferase reporter previously described(15). In brief, 2x10^4^ 293A cells were seeded in each well of a 96 well plate. The next day, cells in each well were transfected with 100 ng PyV NCCR luciferase vectors and 100 ng CMV-renilla plasmid. Luciferase and renilla activity were analyzed two days later using the Dual-Glo Luciferase kit (Promega) following manufacturer’s instructions. The measured luciferase activity in each well was normalized by the corresponding renilla activity.

### Statistics

Statistical analyses were performed using GraphPad Prism 10. Details of individual statistical tests are provided in the relevant figure legends.

## Acknowledgments

This research was funded by NIH/National Cancer Institute R35 grant to D.A.G. (CA209979), a Brave Fellowship awarded by the Brave Like Gabe Foundation to N.J.H.S, and a Fred Hutch Pathogen-Associated Malignancies Integrated Research Center New Technologies pilot award to D.A.G and N.J.H.S.

Assistance with DNA pulldown mass spectrometry experiments was provided by Lisa Jones, Chenwei Lin, and Phil Gafken of the Fred Hutch Proteomics & Metabolomics Shared Resource, which is funded in part through NIH/NCI Cancer Center Support Grant P30 CA015704. The Orbitrap Ascend mass spectrometer used in this research was acquired through the NIH Office of Research Infrastructure Programs grant S10OD030225.

We are very grateful to Dr. Yuan Chang, Dr. Patrick Moore and Dr. Bizunesh Abere (U. Pittsburgh) for sharing their MCPyV minicircle genome with us. We would also like to thank Dr. Jim Pipas and Dr. Sunnie Thompson for insightful discussions at the DNA Tumor Virus Meeting 2025 in Madison, WI, USA.

All cartoons and schematics were created using BioRender.com

