## Supplementary tables for "E2F1-3 activate Merkel cell polyomavirus early transcription and replication"

**Supplementary table 1. sgRNA sequences.**

| Primer name | Sequence (5’ -> 3’) |
|---|---|
| sgNTC | GTGTAGTTCGACCATTCGTG |
| sgF1 | GTATCTAAGGGCAGATCCCA |
| sgF2 | ATAAAAACCACTCCTTAGTG |
| sgR1 | ATACTGCAGTTTCCCGCCCT |
| sgR2 | GCAAATGAGCTACCTCACTA |

**Supplementary table 2. Sequences of 5'-biotinylated primers used to generate DNA pulldown probes**

| dsDNA probe | F primer (5' -> 3') | R primer (5' -> 3') |
|---|---|---|
| AmpR | TAGTGCTGCCATTACCATGAGC | TTACCAGTGCTTGATCAGTGAGG |
| MCPyV NCCR | CCCCATCCTGAAAAATAAATAAG | GACTAAATCCATCTTGTCTATATGC |
| Gorilla PyV NCCR | CCTGAAAAATAAATAAGGATTACTT | GTTGGTGGAGCTCTGCAAGCAAAATG |
| Otomops PyV NCCR | TCTGAAAAAAAAAAAACATGTACTC | GGCTGAAGAGGCTCTAGAGATCCTG |
| Murine PyV (A2) NCCR | TTTGAAAATTCACTTACTTGATCAG | GATGGTGGTGAGGCTGAAATGAGGC |
| Hamster PyV NCCR | AGTTATTAATGAAGTAACTTGGGC | CTTGCTTGTTGCAGCTAGAGATGC |
| TSPyV NCCR | TTTACCTGAAAAATAAGAAAAACTTACC | TTTTGCTGAATGCACCAGAAGACAGG |
| BKPyV (NEB-10) NCCR | GCCTTTGTCCAGTATTAACTGGGG | TTTTGCAAAAAATTGCAAAAGATTAGG |
| JCPyV (CY) NCCR | GGCCAGCTGGTGACAAGCCAAAACAGC | TTTAGCTTTTTGCAGCAAAAAATTAG |
| SV40 NCCR | GGCCTGAAATAACCTCTGAAAG | CTTTGCAAAGCTTTTTGCAAAAGCCTAGG |

**Supplementary table 3. NCCR sequences for pulldown experiments and luciferase vectors.** E2 sites are highlighted and underlined.

| dsDNA probe | NCBI GenBank ID: | Sequence (5' -> 3') |
|---|---|---|
| AmpR | - | TAGTGCTGCCATTACCATGAGCGACAACACCGCGGCCAACT<br>TACTTCTGACAACGATCGGAGGCCCTAAGGAGCTGACTGCA<br>TTTCTTCATAATATGGGTGATCATGTGACCCGGCTTGACCGC<br>TGGGAACCAGAGTTGAACGAAGCCATACCGAACGACGAGC<br>GTGATACCACGATGCCAGTAGCAATGGCCACAACTCTTCGG<br>AAACTACTCACTGGCGAACTTCTTACTCTAGCATCACGACAG<br>CAGCTCATAGACTGGATGGAGGCGGACAAAGTAGCAGGACC<br>ACTTCTTCGCTCGGCCCTCCCTGCTGGCTGGTTCATTGCTG<br>ACAAATCGGGGGCCGGTGAACGCGGCTCTCGCGGCATCATT<br>GCTGCGCTGGGGCCTGATGGTAAGCCCTCACGAATCGTAGT<br>GATCTACACGACGGGGAGTCAGGCCACTATGGACGAACGAA<br>ATAGGCAGATCGCTGAGATCGGTGCCTCACTGATCAAGCACT<br>GGTAA |
| MCPyV NCCR | HM011549.1 | CCTGAAAAATAAATAAGGATACTTACTCTTTTAATGTCCTCCTC<br>CCTTTGTAAGAGAAAAAAAAGCCTCCGGGCCTCCCTTGTTG<br>AAAAAAAGTTAAGAGTCTTCCGTCTCCCTCCCAAACAGAAAG<br>AAAAAAAGTTTTGTTTATCAGTCAAACTCCGCCTCTCCAGGA<br>AATGAGTCAATGCCAGAAACCCTGCAGCAATAAAAGTTCAAT<br>CATGTAACCACAACTTGGCTGCCTAGGTGACTTTTTTTTTTCA<br>AGTTGGCAGAGGCTTGGGGCTCCTAGCCTCCGAGGCCTCT<br>GGAAAAAAAAGAGAGAGGCCTCTGAGGCTTAAGAGGCTTAA<br>TTAGCAAAAAAGGCAGTATCTAAGGGCAGATCCCAAGGGCG<br>GGAAACTGCAGTATAAAAACCACTCCTTAGTGAGGTAGCTCA<br>TTTGCTCCTCTGCTCTTTCTGCAAACTCCTTCTGCATATAGAC<br>AAG |
| Gorilla PyV NCCR | NC_025380.1 | CCTGAAAAATAAATAAGGATTACTTACTCAGCCTTGTCCTCCT<br>CCCTTTGTAAGAGAAAAAAAAGGAGTCTTCTCGCTTCCCTCC<br>TCCCTTTTGAAGAAAAAAAAATGCTGCGTCGCTCTCCCCGCT<br>TGTCGCCTCCCTTTGTGTTGAAAAAAAGTTGTGTTAAGAGTC<br>TACTTCCTCCCTCCCAACTAGATTTAAAAAAATTGTTTATTATAT<br>AACTCCGCCTCTCCAGGATATGAGTCAATGCCAAGAAGCCT<br>GCAGCAATAAAAGTTCAATCAGAGTAAAACCACAAGCTGTCT<br>GCCAGACCACAAGCGTTGCCTAGGCAGCCTATTTTTTTTTAC<br>AAATTAGTGCGAGGCTTGGGGCTCCTAGCCTCCGAGGCCTC<br>TGGAAAAAATAGTGAGAGGCCTCTGAGGCCTCTAACAGCTTA<br>ATTAGCAGAACCATTCCTGGGCGGGAAACTGCAGTATAAAAG<br>CCACTCCTAAGTGATGTAGCTCATTTTGCTTGCAGAGCTCCA<br>CCAAC |
| Otomops PyV NCCR | NC_020071 | TCTGAAAAAAAAAAAACATGTACTCACTTTTAATGCCTCCGCC<br>CGTTCAGAAAGAAAAAAATCCACTCGGCGCTGGGGCTCCCG<br>CCCGCTCTGTTTAAAAAAAATGTTTGAAATGGTTGCTGACCT<br>CCTCCCTTCGTGCTTAGAAAAAATCCTACTCATCATGACTAAC<br>CCCGCCCGCAGAGACAGAAAAAAACAATTTAAAAGGCTGCA<br>GTAAGGAAATGACTCATTCGTGCCGGCGCCTGAACCAAATG<br>ACAGGGGGAGCTCTTTTTTTTTTCAAGTATGCAGAGGCTAGA<br>GGCCCTTAGCCCCTGAGGCTTTCACAGAAAAAGTAGAGAGG<br>CCCTGGGAGGCTTTTTTTAAATTATAGCCGTTAATAGGCGGG<br>AAGGGCTGGTATAAAAGCCTGTTATTCTCCTCCTCAGGATCT<br>CTAGAGCCTCTTCAGCC |

| dsDNA probe | NCBI GenBank ID: | Sequence (5’ -> 3’) |
|---|---|---|
| Murine PyV (A2) NCCR | J02288.1 | TTTGAAAATTCACTTACTTGATCAGCTTCAGAAGATGGCGGA<br>GGGCCTCCAACACAGTAATTTTCCTCCCGACTCTTAAAATAG<br>AAAATGTCAAGTCAGTTAAGCAGGAAGTGACTAACTGACCGC<br>AGCTGGCCGTGCGACATCCTCTTTTAATTAGTTGCTAGGCAA<br>CTGCCCTCCAGAGGGCAGTGTGGTTTTGCAAGAGGAAGCAA<br>AAAGCCTCTCCACCCAGGCCTAGAATGTTTCCACCCAATCAT<br>TACTATGACAACAGCTGTTTTTTTTAGTATTAAGCAGAGGCCG<br>GGGGCCCCTGGCCTCCGCTTACTCTGGAGAAAAAGAAGAG<br>AGGCATTGTAGAGGCTTCCAGAGGCAACTTGTCAAAACAGG<br>ACTGGCGCCTTGGAGGCGCTGTGGGGCCACCCAAATTGATA<br>TAATTAAGCCCCAACCGCCTCTTCCCGCCTCATTTCAGCCTC<br>ACCACCATC |
| Hamster PyV NCCR | NC_001663.2 | AGTTATTAATGAAGTAACTTGGGCATCTATCGCGGCAAAGGC<br>TTCTCCACTAAGTATGGCCTCTACTGAAATTCCAGTAACTGAT<br>GAAATTTCGGAGAGGTAGCTGATCATCTCAATAATCACTGAAA<br>TGGCAGATCCCATGTTGACTTACTTGAACAGTTTGAAAATCTT<br>CTGAACTGTTTCAGGCAGGTTTTTAGGCCGAATTCTAAAGAA<br>ACAGAAAGCAAACACTCAGCGCCGAAGAGCAGGAAATGGCT<br>GACCACTGCACTTGGGCGACACGACACGCCTAGCGATAAGG<br>AAGTCACCATGGCAACATAACCGCAGCACTGCTGTTGTCACA<br>GTTGCCTAGCAAATGACAGACTCAGCAACCACAGGAGAGGA<br>AATGATAGGGCTAGCATTTTTTCAAATGTAAACCAGAGGCTAG<br>GGGCCCCTTGCCTCCTTAGCTCTCAAGTAGAAAAGGAAGAG<br>AGGCTTTTGGGGCTTTTTGGCTTTAAGCCTCATTTTATGAGC<br>AGGAGGAGCTTGTTGCAACTTGAGAGGCGTTTTGAGGCTTC<br>CAGGCAGAGAATACTCACAGACCCCACACAGTCTAGACGCT<br>CAGAAGCATCTCTAGCTGCAACAAGCAAG |
| TSPyV NCCR | NC_014361.1 | TTTACCTGAAAAATAAGAAAAACTTACCATTAGAATTTGAATTT<br>GCCGCGTGAGAACACAGAGCGGGAGGATGTGTTGTTATGGA<br>GACCGGACGGACAGGATGAGAGCAGTTAGACAAGTGCTATC<br>TTAAGAATATACAAAAACAATCACGGGATTTGGCTGGCTTCCT<br>CTTGTCTCTTGGCTGCCTTCCTGTGTTTGTGGCAGTATGACA<br>AACATCCCCTTGAGACTAACTGACACAGCATTTTCCATAACAA<br>ATGATAGGTGGCGCACGAGCTTCCTCAGGATAACGGTCTTAA<br>GCAAACAATGACCTATTTATGTCAACACTCTAAGTATAGTTCC<br>TCAAATTCCTAATAGGGGGTTACTATGGAAATCTTGTTTTTTCT<br>CAGTATTACCAGGAGGCTGAGGCTTCCTGCCCCCTCTTGAC<br>ACAGACATAGAGGGAGAGGCCTCCGGAGGCCTCTTGAGGC<br>TTGGCACTTCCTCCCTTTTATGTAAGCCAAATCTGATGAAAAT<br>TAATTGAGGCACACAGAGGCTTTAAAAGGGCTCCAAAAAGCT<br>CCCATCTTCATTTCTTCATTTCTCCCCTTACCTGTCTTCTGGT<br>GCATTCAGCAAA |
| BKPyV (NEB-10) NCCR | AB365141.1 | GCCTTTGTCCAGTATTAACTGGGGACAAGGCCAAGATTCCTA<br>GGCTCGCAAAACATGTCTGTCTGGCTGCTTTCCACTCCTTTG<br>GCTAGTTTCCACTTCCTCTTGTGTTTATTTGAGAATTCTAGGG<br>GCGGGGTTTCACTATTAACTGCCACTGGCTGGCTGCCCAGT<br>CATGCACTTTCCTTCCTGAGGTCATGGTTTGGCTGCATTCCA<br>TGGGAAAGCAGCTCCTCCCTGTGGCCTTTTTTTTTATAATATA<br>TAAGAGGCCGAGGCCGCCTCTGCCTCCACCCTTTCTCTCAA<br>GTAGTAAGGGTGTGGAGGCTTTTTCTGAGGCCTAGCAAAAC<br>TATTTGGGGAAATCCCTAATCTTTTGCAATTTTTTGCAAAA |

| dsDNA probe | NCBI GenBank ID: | Sequence (5’ -> 3’) |
|---|---|---|
| JCPyV (CY) NCCR | AB038249.1 | GGCCAGCTGGTGACAAGCCAAAACAGCTCTGGCTCGCAAAA<br>CATGTTCCCCTGGCTGCTTTCCACTTCCCCTTGTGCTTTGTT<br>TACTTGTGATTAAGGACTATGGGAGGGGTTTCACTATAACTGC<br>CAGTGGCATGCAGCCAGGGCTCCCTCTGGCTGTCAGCTGG<br>TTGGCTCCCTAGGTATGAGCTCATGCTTGGCTGGCAGCCATC<br>CAGTTTTAGCCAGCTCCTCCCTACCTTCCCTTTTTTTTATATAT<br>ACAGGAGGCCGAGGCCGCCTCCGCCTCCAAGCTTACTCAG<br>AAGTAGTAAGGGCGTGGAGGCTTTTTAGGAGGCCAGGGAAA<br>TTCCCTTGTTTTTCCCTTTTTTGCACTAATTTTTTGCTGCAAAA<br>AGCTAAA |
| SV40 NCCR | J02400.1 | GGCCTGAAATAACCTCTGAAAGAGGAACTTGGTTAGGTACCT<br>TCTGAGGCGGAAAGAACCAGCTGTGGAATGTGTGTCAGTTA<br>GGGTGTGGAAAGTCCCCAGGCTCCCCAGCAGGCAGAAGTA<br>TGCAAAGCATGCATCTCAATTAGTCAGCAACCAGGTGTGGAA<br>AGTCCCCAGGCTCCCCAGCAGGCAGAAGTATGCAAAGCATG<br>CATCTCAATTAGTCAGCAACCATAGTCCCGCCCCTAACTCCG<br>CCCATCCCGCCCCTAACTCCGCCCAGTTCCGCCCATTCTCC<br>GCCCCATGGCTGACTAATTTTTTTTATTTATGCAGAGGCCGAG<br>GCCGCCTCGGCCTCTGAGCTATTCCAGAAGTAGTGAGGAGG<br>CTTTTTTGGAGGCCTAGGCTTTTGCAAAAAGCTTTGCAAAG |

**Supplementary table 4. qPCR primers.**

| **Primer name** | **Sequence (5' -> 3')** |
|---|---|
| PanT-7 F | ATGGCAACATCCCTCTGATGA |
| PanT-7 R | TGGAATTTGCTCCAAAGGGTG |
| 36B4 F | TGCCAGTGTCTGTCTGCAGA |
| 36B4 R | ACAAAGGCAGATGGATCAGC |
| VP1 F | AAAACACCCAAAAGGCAATG |
| VP1 R | GCAGAGACACTCTTGCCACA |
| GAPDH F | TGTGTCCCTCAATATGGTCCTGTC |
| GAPDH R | ATGGTGGTGAAGACGCCAGT |

**Supplementary dataset 1. MKL-2 DNA pulldown mass spectrometry dataset**. Fold change (FC) and p values for proteins detected by mass spectrometry following AmpR and NCCR pulldowns. $\log_2$(FC), and $-\log_{10}$(p) are plotted in Figure 1B.

**Supplementary dataset 2. 293A DNA pulldown mass spectrometry dataset**. Fold change (FC) and p values for proteins detected by mass spectrometry following AmpR and NCCR pulldowns. $\log_2$(FC), and $-\log_{10}$(p) are plotted in Figure 2A.
